# A filter-flow perspective of hematogenous metastasis offers a non-genetic paradigm for personalized cancer therapy

**DOI:** 10.1101/000125

**Authors:** Jacob G. Scott, Alexander G. Fletcher, Philip K. Maini, Alexander R. A. Anderson, Philip Gerlee

**Affiliations:** Integrated Mathematical Oncology, H. Lee Moffitt Cancer Center and Research Institute, Tampa, FL, USA; Wolfson Centre for Mathematical Biology, Mathematical Institute, Radcliffe Observatory Quarter, Oxford University, Woodstock Road, Oxford OX2 6GG, UK; Mathematical Sciences, University of Gothenburg and Chalmers University of Technology, SE-41296 Gothenburg, Sweden

**Keywords:** metastasis, mathematical model, cancer, personalized medicine, oligometastasi

## Abstract

**Translational Relevance:** Since the discovery of circulating tumor cells (CTC), we have struggled for ways to use them to inform treatment. The only currently accepted method for this is a ‘more is worse’ paradigm by which clinicians measure CTC burden before and after treatment to assess efficacy. Research efforts are currently focused almost entirely on genetic classification of these cells, which has yet to bear any fruit translationally. We suggest that we should shift the focus of our investigation to one driven by a physical sciences perspective. Specifically, by understanding the vascular system as a network of interconnected organs and capillary beds as filters that capture CTCs. By ascertaining the distribution of CTCs in this network for individual patients, information about the existence of subclinical metastatic disease, and therefore metastatic propensity, will come to light, and allow for better staging, prognostication and rational use of organ-directed therapy in the setting of oligometastatic disease.

**Abstract:** *Purpose:* Research into mechanisms of hematogenous metastasis has largely become genetic in focus, attempting to understand the molecular basis of ‘seed-soil’ relationships. However, preceding this biological mechanism is the physical process of dissemination of circulating tumour cells (CTCs) in the circulatory network. We utilize a novel, network perspective of hematogenous metastasis and a large dataset on metastatic patterns to shed new light on this process.

*Experimental Design:* The metastatic efficiency index (MEI), previously suggested by Weiss, quantifies the process of hematogenous metastasis by taking the ratio of metastatic incidence for a given primary-target organ pair and the relative blood flow between the two sites. In this paper we extend the methodology by taking into account the reduction in CTC number that occurs in capillary beds and a novel network model of CTC flow.

*Results:* By applying this model to a dataset of metastatic incidence, we show that the MEI depends strongly on the assumptions of micrometastatic lesions in the lung and liver. Utilizing this framework we can represent different configurations of metastatic disease and offer a rational method for identifying patients with oligometastatic disease for inclusion in future trials.

*Conclusions:* We show that our understanding of the dynamics of CTC flow is significantly lacking, and that this specifically precludes our ability to predict metastatic patterns in individual patients. Our formalism suggests an opportunity to go a step further in metastatic disease characterization by including the distribution of CTCs at staging, offering a rational method of trial design for oligometastatic disease.

## Introduction

Nearly 150 years after Ashworth’s discovery of the circulating tumor cell (CTC) (1), the putative vector of hematogenous metastatic disease, the mechanisms driving this process remain poorly understood and unstoppable (2). For over a century the dominant paradigm has been the seminal, yet qualitative, seed-soil hypothesis proposed by Paget in 1889 (3). This began to be challenged in 1992, when a quantification of the contribution of mechanical and seed-soil effects was attempted by Weiss (4), who considered the ‘metastatic efficiency index’ (MEI) of individual primary tumors and metastatic sites (5). He calculated MEI as the ratio of metastatic involvement to blood flow through an organ and three classes of organ pairs emerged: low, where the soil-organ relationship is hostile; high, where it is friendly; and medium, where blood flow patterns to a large extent explain patterns of metastatic spread. The utility of Weiss’ classification method largely ended there, and has since been put aside in favor of genetic investigations (6, 7), an exception being work in prostate cancer by Pienta and Loberg (8) showing a lack of correlation between blood flow and incidence, suggesting strong seed-soil effects. While illuminating, the gene-centric approach has yet to offer any actionable conclusions, and its applicability is threatened by the growing understanding that genetic heterogeneity, not clonality, is the rule in cancer (9, 10, 11). Our aim is to revisit the pre-genetic model and show that a physical perspective of metastatic spread can lead to new and actionable insights into this enigmatic disease process.

While primary tumour and lymph node metastases are carefully described in the clinic, metastatic disease is considered to be a binary change of state, a patient being diagnosed with either M0 or M1. Until recently, this was appropriate, as even perfect information about the existence and distribution of metastatic disease would have done little to affect treatment choice, the options being limited to the use of systemic chemotherapies. However, recent years have witnessed the advent of more effective and tolerable localized therapies for metastatic involvement, in the form of liver-directed therapy (12), bone-seeking radionuclides (13) and stereotactic body radiation therapy (14, 15). These recently adopted modalities have allowed for targeted therapy to specific parts of the body with minimal side efffects and high eradication potential. Further, trials offering treatment with curative intent to patients with limited, ‘oligometastatic’ disease have shown promise (14, 15), although it is not yet possible to identify such patients in an objective manner (16, 17). The time is therefore ripe for a quantiative framework that can analyse and guide these and similar efforts.

In this paper we seek to use Weiss’ MEI and a recently proposed model of CTC dynamics (summarised in Figure 1) (18, 19) to extend our understanding of metastasis. This framework presents a way to utilize ‘personalized’ patient CTC measurements to assay for the burden and distribution of metastatic disease. These measurements represent a *novel class of patient-specific data* by which any pattern of metastasis can be understood. This allows for a new way to dissect out the heterogeneous groups from population level data, and hence represents a non-genetic, translatable method by which to alter staging and hence treatment strategies.

**Figure 1:**
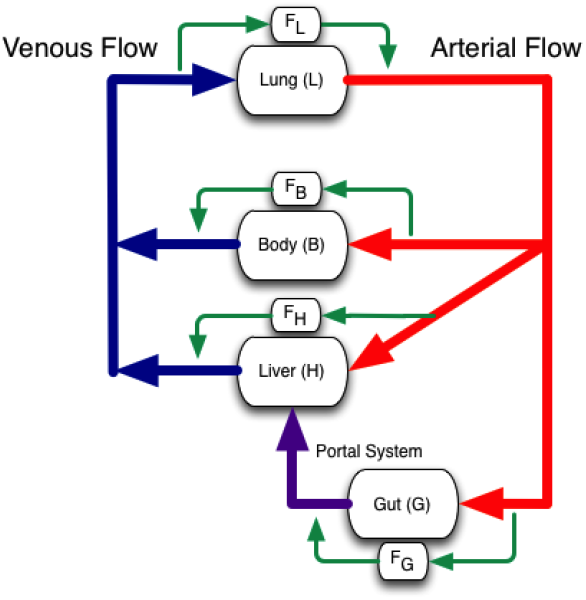
Schematic of the human vascular system network topology. It is evident by inspection of the network diagram that tumors originating in the gut and lung experience significantly different flow patterns and a different order in which they experience filtration at capillary beds than tumors originating in other parts of the ‘body’ (18). The alternate pathways (green) represent the fraction of cells which evade arrest (filtration) at a given capillary bed. There are scant measurements of this fraction in the literature, and none in clinical studies that evaluate outcomes.

To do this, we consider blood flow between organs (20), filtration in capillary beds (see Figure 1) and distribution of metastatic involvement in a series of untreated patients at autopsy (21). For each organ-organ pair we calculate the MEI by normalising incidence by putative CTC flow between the two organs, taking into account the reduction that occurs in capillary beds (18, 19). This post-capillary bed reduction in CTC numbers can be altered by the presence of micrometastases, which can amplify CTC numbers downstream of their location through shedding. Thus, by adjusting filtration rates throughout the network, we can represent different configurations of metastatic disease and hence capture different organ-organ metastatic efficiencies.

## Materials and Methods

### Calculation of Metastatic Efficiency Index (MEI)

The autopsy dataset used in the analysis covers 3827 patients presenting with primary tumors in 30 different anatomical sites (21). For each primary tumor the number of metastases are reported according to anatomical site (in total 9484 metastases). As we focus on the effect of blood flow patterns, we consider only the organs for which blood flow has been measured.

For each organ-organ pair we calculate the metastatic involvement *N*_*ij*_ as the ratio between the number of occurrences of the primary tumor in organ *i* and the number of metastases reported in the target organ *j*, for each pair (*i*, *j*) presented in the autopsy data (21). We have that 0 *≤ N*_*ij*_ *<* 1 and this number corresponds to the fraction of cases where a primary tumor in organ *i* gave rise to a metastasis in organ *j*. The metastatic efficiency index (MEI) from organ *i* to *j* is then defined by *M*_*ij*_ = *N*_*ij*_/*ϕ*_*ij*_, where *ϕ*_*ij*_ is the relative flow of CTCs from organ *i* to *j*. This quantity takes into account the blood flow that each target organ receives and the reduction in CTCs that occurs en route between the two organs. For the sake of simplicity we consider only the effects of capillary bed passage, and it has been shown in clinical studies that approximately 1 in 10 000 CTCs remain viable after such a passage (22) and in animal studies that these cells are able to initiate tumours (23). We thus assume that there occurs a reduction of CTC number by a factor *F*_*k*_ when the cells pass through organ *k*. As a baseline, we use the pass rate *F*_*k*_ = 10^*−*4^ for all organs.

It is well known that metastases in the lung and liver have the ability to shed cells into the bloodstream and hence give rise to ‘second order’ metastases (24), and it has been shown that even ‘dormant’ micrometastatic disease can shed CTCs (25). If one were to measure the CTC concentration downstream of an organ containing metastases, then it would be higher than in the case of a disease-free organ. For our purposes, this implies that the presence of metastatic disease can be represented in the model as a lower reduction of CTCs in the capillary bed of the affected organ. This simplification is only valid if we disregard the biological properties of the CTCs (since CTCs originating from metastases might have different genotypes and phenotypes compared to cells from the primary tumor), but is sufficient for our purposes. To simulate the presence of micrometastases in the lung and liver we therefore change the pass rates to *F*_*L*_ = 10^*−*1^ and *F*_*H*_ = 10^*−*1^ respectively.

As an example of our methodology, we now present the calculations for the MEI for breast to adrenal gland. The cancer cells leaving a breast tumor enter the circulation on the venous side and are transported via the heart to the lung capillary bed, through which only a fraction *F*_*L*_ pass as viable cells. These cells then flow into the arterial side of the circulation and are randomly distributed to the different organs of the body according to blood flow, of which the adrenal gland receives 0.3% (20). The relative flow of CTCs from breast to adrenal gland is therefore given by *ϕ*_*breast*,*adrenal*_ = *F*_*L*_ × 0.3 = 0.3 × 10^*−*3^.

To illustrate the effect of micrometastatic disease on MEIs, we have compared Weiss’ original method with MEIs calculated using the filter-flow framework in four different regimes: no micrometastases, micrometastases present in the lung, in the liver or in both locations. Figure 2 shows the result of this comparison for four organ pairs. We see that Weiss’ metric differs from ours, but more importantly that the metastatic efficiency depends on the current disease state. For example, our estimate of the efficiency with which cells orginating from a primary pancreatic tumor can form kidney metastases varies over six orders of magnitude, depending on whether micrometastatic lesions are present, and their location. This effect highlights an opportunity to go a step further in disease characterization than presence or absence of CTCs at staging - we need to include information about where in the vascular network, and in what relative quantities, these CTCs reside.

**Figure 2:**
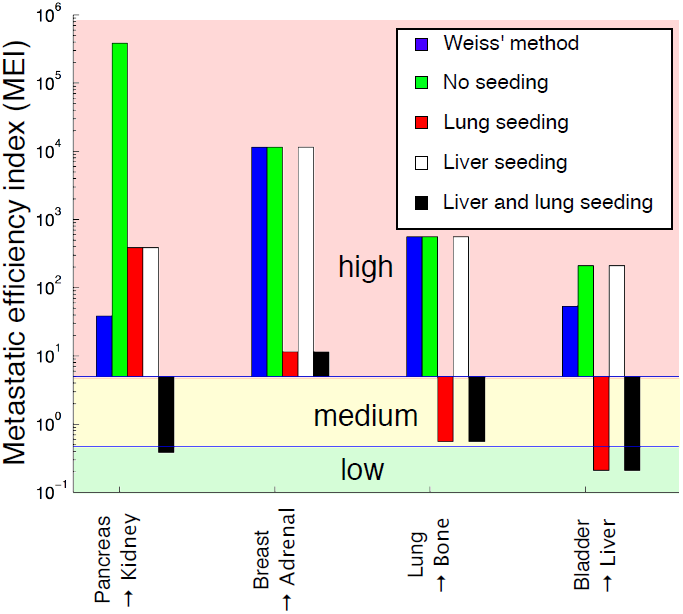
The impact of filter and flow characteristics on estimation of the metastatic efficiency index (MEI). We have compared Weiss’ original method with our filter-flow framework under the assumption of no micrometastases, micrometastases in the lung, in the liver, and in both locations. The comparison is carried out for four organ pairs that cover the canonical pathways of spread (gut *→* body, body *→* body, lung *→* body and body *→* liver). We see that because Weiss’ method only considers the dynamics on the arterial side it underestimates the MEI in two of the cases (pancreas *→* kidney, and bladder *→* liver). From the comparison it is also evident that assumptions about the presence or absence of micrometastases heavily influences the results, in the case of pancreas *→* kidney shifting the MEI six orders of magnitude from a low MEI to a high one (as defined by Weiss (5)).

### Patient group decomposition

The preceding analysis assumes that the filtration rate for each organ is identical for all patients in the data set. This is likely a gross oversimplification, but no clinical trial has yet determined the intrapatient heterogeneity in this (currently absent) parameter set. Previously we used incidence data to calculate MEIs, but we may also reverse the process and calculate the prevalence of micrometastatic disease given incidence data and organ-pair MEIs. We now show how this can be used as a means to suggest possible patient group decompositions.

The incidence, *N*_*ij*_, relative flow of CTCs, *ϕ*_*ij*_ and the MEI, *M*_*ij*_ are related according to *M*_*ij*_ = *N*_*ij*_/*ϕ*_*ij*_, or equivalently *ϕ*_*ij*_ = *N*_*ij*_/*M*_*ij*_. We now assume that the MEI is fixed, while the flow *ϕ*_*ij*_ varies across patients, and consider four patient groups: no micrometastases, micrometastases in the lung, micrometastases in the liver and micrometastases in both. If we now let *n*_*k*_ denote the fraction of patients in each group, *k*, where ∑*n*_*k*_ = 1, then we can write

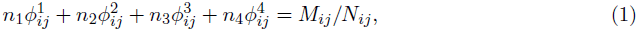

where 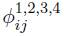 is the relative flow of CTCs in the different patient groups. The problem of finding the *n*_*k*_’s is underdetermined, and the solution is given by any point on a surface defined by (1), such that *n*_*k*_ *>* 0 for all patient groups and ∑_*k*_*n*_*k*_ = 1. This implies that aggregated incidence data can be explained by many different patient group compositions, each with its distinct pattern of metastatic progression. Let us illustrate this with a concrete example based on the autopsy dataset utilized above.

The population incidence of metastases in the adrenal gland arising from primaries in the large intestine equals 7.5%, and by fixing the MEI and using the above method, the incidence rate can be explained by a subdivision according to 25% in the no metastasis group, 20% in the liver metastases group, 5% in the lung metastases group, and 50% of the patients harboring metastases in both liver and lung. However the incidence can also be explained by a subdivision of 5%, 25%, 20% and 50% into each patient group respectively. This highlights the fact that population-based measures of incidence cannot be used to predict individual patient metastasis dynamics as different patients can exhibit fundamentally different patterns of spread.

## Discussion

To enable these insights and their translation to the clinic, systematic testing of individual patient filter-flow parameters is required. While several groups have successfully interrogated this step of the metastatic cascade in animals (26, 27, 28), it has yet to be done in humans. To effect this, measurement of CTCs from each of the individual vascular compartments (see Figure 3) at initial staging and subsequent correlation with outcomes would yield initial information with which more complete models could be built, and from which rational prospective trials could be designed. This level of understanding of an individual patient’s disease state constitutes a new type of personalized medicine, which seeks to assay not just the collection of mutations that a patient’s cancer cells have accumulated, but also their physical distribution. This would allow for more accurate staging and the rational inclusion of organ directed therapy in clinical trials, a concept which is gaining popularity with recently approved methods existing for bone and liver (13, 12). We present an example of how this methodology could be implemented in Figure 3, in which patients presenting with stage II colorectal cancer are stratified after resection of their primary tumour according to CTC burden in different compartments of the circulatory system. We have chosen stage II colon cancer as a first approach as there are no clear guidelines (29) for which patients should get adjuvant therapy, and further because we have effective liver-directed therapies which could be used in the adjuvant setting. In this case, our methodology would offer a rational method of treatment allocation - offering a way to spare patients from systemic therapy and its risks. While we have chosen to highlight colorectal cancer, we stress that this sort of approach, and the information gleaned from it, would be useful for any primary cancer.

**Figure 3:**
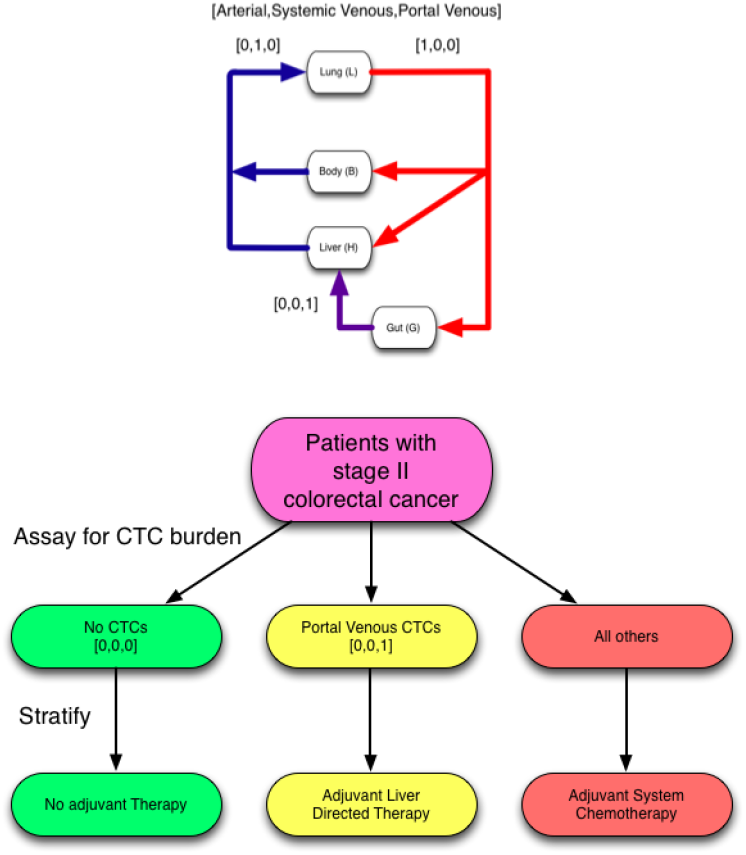
An example of clinical trial stratification based on CTC localization. Stage II colon cancer remains enigmatic, with no clear guidelines for adjuvant therapy after surgery (29). We propose that stratifying by CTC presence or absence in specific vascular compartments, information about subclinical metastatic disease could be brought to light, and recommendations for location specific treatment, if appropriate, could be made. A first approximation would be to collect this information prospectively in the setting of existing trials.

## Conclusion

We have presented a fresh look at old data on metastatic patterns inspired by a physical science perspective and shown that there is a deep gap in our understanding. Specifically, we show that our lack of knowledge of the interaction dynamics of CTCs in foreign organ capillary beds prevents us from making further progress towards predicting patterns of spread. We suggest some simple steps to fill in these gaps, and a simple trial design to take advantage of some of currently obtainable, yet overlooked, patient specific information: CTC distribution throughout the patient’s vascular network.

By further elucidating the principles underlying hematogeonous metastasis, we hope to make inroads toward therapeutic strategy changes that would otherwise be impossible. Our results highlight the value of the physical perspective of the metastatic cascade, and the importance of addressing not only genetic factors, but also physiological and anatomical aspects of the process, which in this gene-centric era have been largely forgotten.

